# Cytokine expression profile in the human brain of older adults

**DOI:** 10.1101/2025.06.02.657444

**Authors:** Juliana Beker Godoy, Ricardo A. Vialle, Loren dos Santos, Roberto T Raittz, Yanling Wang, Vilas Menon, Philip L. De Jager, Julie A. Schneider, Shinya Tasaki, David A. Bennett, Dieval Guizelini, Katia de Paiva Lopes

## Abstract

Alzheimer’s disease (AD) is a complex neurodegenerative condition linked to chronic neuroinflammation. This study investigates the cytokine gene expression profile in cortical tissue samples from elderly individuals with and without AD to identify potential biomarkers and enhance our understanding of disease pathogenesis. Utilizing high-depth RNA sequencing data, we identified a set of cytokines whose expression significantly associated with different aspects of the AD phenotype, including measures of neurofibrillary tangles, amyloid-β deposition, and a person-specific rate of cognitive decline. Single-nucleus transcriptomics data facilitated the identification of specific cell types, such as microglia and oligodendrocytes, that significantly contribute to the inflammatory response in AD. Additionally, we observed a strong correlation between the expression of certain cytokines and genetic risk for the disease. Our findings indicate that cytokine-mediated neuroinflammation plays a vital role in AD progression and that modulating the immune response may offer a promising strategy for developing new therapies.

## INTRODUCTION

Alzheimer’s disease (AD) is the most common cause of dementia and one of the top ten causes of death worldwide. Even though it is expected to affect 150 million people by 2050, no effective treatment or diagnosis is currently available (Alzheimer’s Association 2024; Naghavi et al. 2024). AD is a neurodegenerative disorder characterized by extracellular β-amyloid plaques and intracellular hyperphosphorylated tau tangles (Long and Holtzman 2019). While it typically manifests with memory loss, it can also affect speech, visuospatial skills, and executive functions (Knopman et al. 2021). Cognitive impairment in AD varies, with early signs often appearing as subjective decline despite normal assessments. Mild cognitive impairment (MCI) marks the initial symptomatic stage, featuring slight deficits in one or more cognitive domains while maintaining daily functioning (Assaf and Tanielian 2018). Recent anti-amyloid antibody trials have challenged the notion that β-amyloid accumulation is the sole cause of Alzheimer’s disease. The findings indicate a more complex disease process involving various interacting factors, where amyloid pathology may act as a necessary trigger but is insufficient to drive the disease fully (Kepp et al. 2023; Selkoe and Hardy 2016).

For many years, neuroinflammation was considered a secondary response to neuronal death. However, a great number of epidemiological, clinical, and experimental studies now widely recognize neuroinflammation as a central feature of AD, highlighting its key role in the disease’s pathological process (Heneka et al. 2015; Prins et al. 2022; Castro-Gomez and Heneka 2024). Genome-wide association studies (GWAS) have shown that over 51% of Alzheimer’s disease risk genes are enriched in innate immune system pathways (Bellenguez et al. 2022; Kunkle et al. 2019; Jansen et al. 2019). Additionally, rare AD risk variants have been identified in genes crucial for immune responses, particularly those involved in microglial activation and phagocytic functions (e.g *TREM2*, *APOE*) (Ray et al. 2024; Bis et al. 2020; Keren-Shaul et al. 2017).

Building on the understanding that the immune system pathways are crucial to AD risk (Raj et al. 2014), neuroinflammation has emerged as a central aspect of the disease’s progression. Neuroinflammation is initiated by activating the brain’s innate immune system in response to protein buildup, with cytokines acting as key mediators (Sobue, Komine, and Yamanaka 2023). Produced by activated glial cells and neurons, these small signaling molecules include interleukins, chemokines, tumor necrosis factors, interferons, and growth factors. Under healthy conditions, cytokines are crucial to neurodevelopment and regulating cognitive functions like synaptic neurotransmission, brain plasticity, learning, and memory retention (Bourgognon and Cavanagh 2020; Zipp, Bittner, and Schafer 2023). On the other hand, cytokines promote inflammation through intercellular communication, accelerating AD progression (Chen et al. 2024). Previous studies have shown that elderly individuals with β-amyloid neuropathology and cognitive decline exhibit increased brain cytokine expression (Flores-Aguilar et al. 2021). Here, we extend these findings by leveraging data from two longitudinal cohorts, the Religious Orders Study and the Rush Memory and Aging Project (ROS/MAP) (David A. Bennett et al. 2018) (**Figure 01**). We investigated cytokine expression in the cortical brain tissue of community-dwelling older adults and its role in AD and related disorders (ADRD). First, we examined the association of these genes with AD pathology and cognitive decline. Next, we explored cell-type-specific expression profiles using single-nucleus RNA sequencing (snRNA-Seq). Then, we evaluated the relationship between AD genetic risk and cytokine expression. Finally, we conducted cell communication analysis to understand the intercellular communication mediated by these inflammatory molecules.

**Figure 1:**
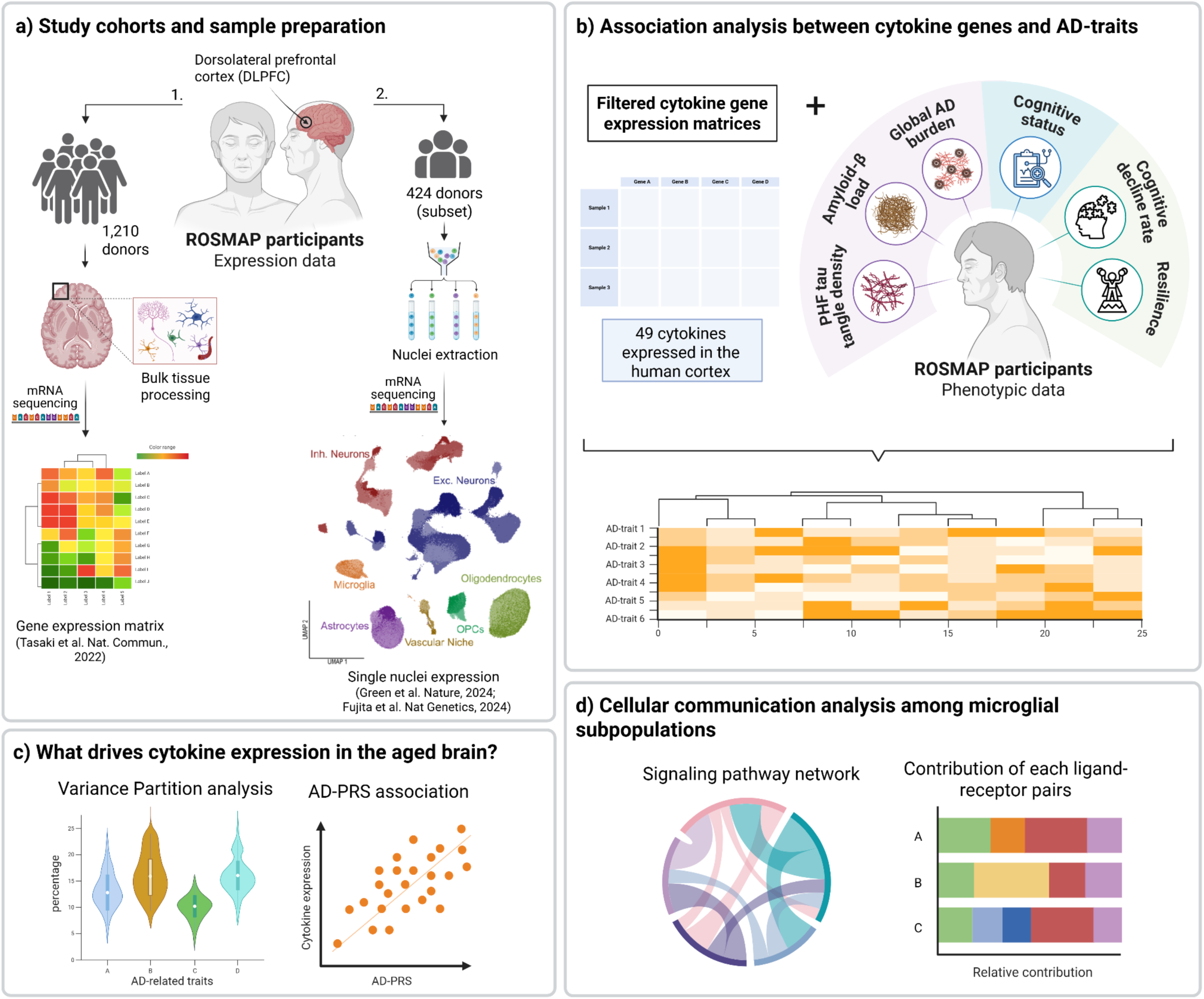
Study design overview. **a)** This study analyzed (1) bulk RNA-Seq data (n = 1,210) and a subset of (2) snRNA-Seq data (n = 424) from the DLPFC region of ROSMAP participants. Figure created with Biorender. **b)** Expression matrices were filtered to retain only cytokine genes, and their associations with selected AD traits were tested using linear and/or logistic regression models. **c)** VariancePartition analysis and AD genetic risk scores (AD-PRS) were used to identify potential expression drivers. **d)** Cell-cell communication analysis among microglial subpopulations revealed cytokine signaling pathways in human-aged brains.

## RESULTS

### Characteristics of the participants

Transcriptomic data originate from participants of two longitudinal clinical-pathological studies of aging and dementia, the Religious Orders Study (ROS) and the Rush Memory and Aging Project (MAP), commonly referred to as ROSMAP. Both studies were approved by the Institutional Review Board of Rush University Medical Center (Chicago, IL). The participants are older adults who enroll without dementia and agree to longitudinal clinical evaluations and organ donation. To date, ROSMAP has enrolled over 4,000 participants and conducted more than 2000 brain autopsies. The bulk RNA-Seq data from the DLPFC were available for 1,210 participants (**Supplementary Table 01-02**), and the snRNA-Seq for a subset of 424 participants. The participants enrolled at a mean age of 80.8 (SD: 7.0) years and died at an average age of 89.5 (SD: 6.6), with a third being female (∼70%). At death, 32.9% had no cognitive impairment, 23.6% had mild cognitive impairment (MCI), and the remaining 43.5% had dementia. At autopsy, 64% had a pathologic AD diagnosis. The characteristics were similar for the participants with snRNA-Seq data (**Supplementary Table 03**).

### Cytokine expression associated with AD phenotype in specific brain cell types

First, we investigated the associations between cytokine expression from bulk RNA-Seq data and seven AD-related traits. These traits included the accumulation of AD pathology measured by PHFtau tangle density and amyloid-β load, global AD pathology capturing the overall burden of AD in the aged brain, Alzheimer’s dementia diagnosis, MCI diagnosis, the rate of cognitive decline for each participant, and a measure of cognitive resilience. Overall, 20 out of 42 cytokines (47.6%) were associated with at least one of the tested traits (adj. *p*-value < 0.05) (**Figure 2a** and **Supplementary Table 04**). Specifically, 7 cytokines were associated with amyloid-β load; 9 with MCI diagnosis; 10 with global AD burden and resilience; 14 with PHFtau tangle density and AD diagnosis; and 15 with cognitive decline. The strongest association was observed between *IL15* expression and cognitive decline (adj. *p*-value = 9.79 × 10^−10^), as highlighted in **Figure 2b**. As expected, these associations overlapped across traits. For example, 12 cytokines (*DCN*, *IGF1*, *HGF*, *THPO*, *TGFB1*, *CCL2, TNFSF4*, *SPP1*, *IL33*, *CTF1, IL15*, and *CSF1*) were significantly associated with both cognitive decline and AD pathology, while 4 cytokines (*TGFB1*, *SPP1*, *CTF1*, and *CSF1*) were associated with all the tested traits. For replication, we expanded our analysis to external bulk RNA-Seq datasets from the Mayo Clinic and Mount Sinai Brain Bank to assess generalizability. 34 out of 44 cytokines (77.3%) were associated with Thal, Braak, or AD diagnosis in the Mayo bulk RNA-Seq. Also, 15 out of 47 (31.9%) cytokines were associated with Braak or CDR in the Mount Sinai data (**Supplementary Figures 1a-b** and **Supplementary Tables 05-06**).

**Fig 2:**
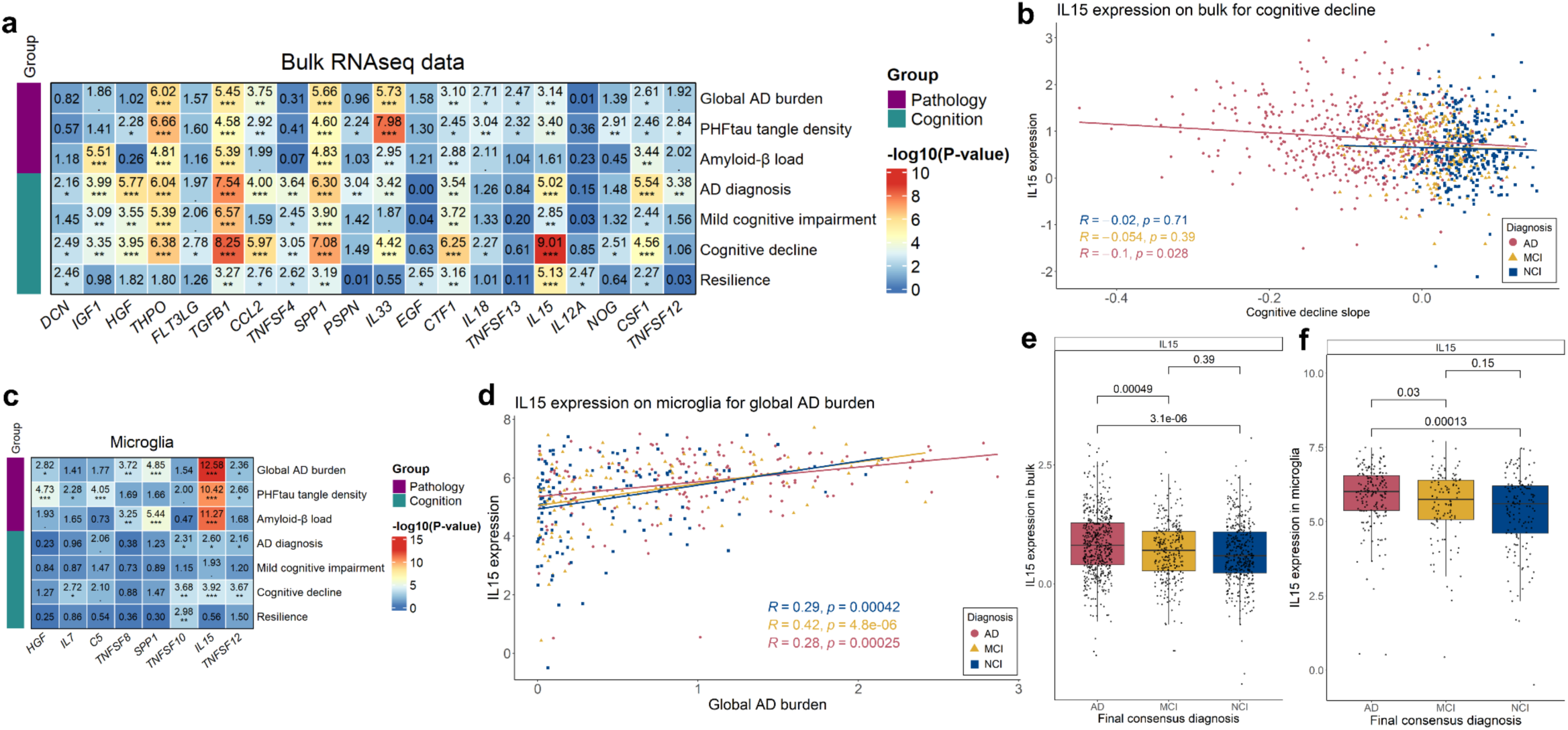
Association analysis between brain inflammatory cytokine expression and AD traits. **a)** Heatmap illustrating the significant associations between cytokine expression levels and AD phenotypes, represented by -log_10_(*p*-value) (***FDR *p*-value < 0.001, **FDR *p*-value < 0.01, *FDR *p*-value < 0.05). Linear and/or logistic regressions were performed to explore the relationships between cytokine expression and phenotypic data. The AD-traits are labelled according to their category: Pathology (global AD pathology burden, PHFtau tangle density, and amyloid-β load) and Cognition (AD clinical diagnosis, MCI diagnosis, cognitive decline, and resilience). All models were adjusted for age, sex, and years of education to control for potential confounding factors. In the bulk data, 20 cytokines were associated with phenotypic data. **b)** Scatter plot showing the association of *IL15* expression in the bulk data with global AD burden. Participants were categorized based on their clinical diagnosis (AD, MCI, or NCI), with each dot color-coded accordingly. **c)** Heatmap of significant associations between cytokine expression levels in microglia from the DLPFC region (columns) and phenotypic data (rows). Colors represent the -log_10_ of the *p*-value, ranging from highest (red) to lowest (blue). **d)** Scatter plot illustrating the association between *IL15* expression in microglia and cognitive decline. Participants were categorized by clinical diagnosis (AD, MCI, or NCI), with each dot color-coded to reflect their group. **e)** Differential expression level of the *IL15* in bulk data for individuals diagnosed with Alzheimer’s dementia (AD; red), mild cognitive impairment (MCI; yellow), and no cognitive impairment (NCI; blue). Data are presented in boxplots, with the median indicated by a horizontal line. Nominal *p*-values are shown at the top and calculated using one-sided t-tests. **f)** Differential *IL15* expression in microglia across AD (red), MCI (yellow), and NC (blue). Data are shown in boxplots with medians as horizontal lines. Nominal *p*-values, calculated using one-sided t-tests, are displayed at the top.

Because cytokine production levels vary among different cell types, we investigated their associations with the same AD traits across the seven major cell types in the DLPFC. Cytokines were expressed in all cell types and significantly associated with AD traits, except endothelial cells (**Supplementary Figure 2** and **Supplementary Table 7**). We identified 4 cytokines associated with AD traits in oligodendrocytes, 6 in inhibitory neurons and OPCs, 8 in microglia, 9 in excitatory neurons, and 10 in astrocytes. A comparison of the datasets reveals that *SPP1* levels were significantly associated with all AD tested traits in bulk data but exhibited distinct patterns across cell types. In oligodendrocytes, *SPP1* levels were strongly associated with global AD burden (adj. *p*-value = 5.92 × 10^−4^), PHFtau tangle density (adj. *p*-value = 3.99 × 10^−6^), amyloid-β load (adj. *p*-value = 1.35 × 10^−2^), cognitive decline (adj. *p*-value = 4.36 × 10^− 6^), and resilience (adj. *p*-value = 4.12 × 10^−2^), similar to bulk analysis (**Supplementary Fig. 2d**). However, *SPP1* expression in microglia was associated only with global AD burden (adj. *p*-value = 9.98 × 10^−5^) and amyloid-β load (adj. *p*-value = 2.51 × 10^−5^) (**Figure 2c**). Notably, the strongest association was also observed in microglia, between global pathological burden and *IL15* levels (adj. *p*-value = 1.83 × 10^−12^; **Figure 2c**). Consistently, *IL15* overexpression was positively correlated with higher pathology deposition in MCI participants (R = 0.42; *p*-value = 4.8 × 10^−6^; **Figure 2d**). We also observed a strong association between *IL15* expression and PHFtau tangle density (adj. *p*-value = 2.64 × 10^−10^) and amyloid-β load (adj. *p*-value = 3.72 × 10^−11^) in this cell type. Interestingly, this last association was observed exclusively in microglia and was not generalizable to the bulk data. Both bulk and snRNA-Seq data showed significantly upregulated *IL15* levels in AD participants compared with MCI (bulk *p*-value = 0.00049; snRNA-Seq *p*-value = 0.03) and NCI (bulk *p*-value = 3.1 × 10^−6^; snRNA-Seq *p*-value = 0.00013; **Figures 2e-f**).

Some cytokine genes were detected exclusively in the snRNA-Seq data and exhibited distinct association patterns. The expression of the pro-inflammatory cytokine *IL7* was associated with PHFtau tangle density and amyloid-β load in OPCs (adj. *p*-value < 0.05; ; **Supplementary Figure 2e**), amyloid-β and cognitive decline in inhibitory neurons (adj. *p*-value < 0.05; ; **Supplementary Figure 2e**), and only with amyloid-β in excitatory neurons (adj. *p*-value ≤ 0.001; **Supplementary Figure 2a**). On the other hand, *TNFSF8* was significantly associated with AD traits only in microglia, particularly with global AD burden and amyloid-β load (adj. *p*-value < 0.005; **Figure 2c**). In summary, these findings reveal distinct neuroinflammation patterns across different brain cell types, with varying neuropathologic associations suggesting that each cell type has a unique relationship with AD.

### Biological factors driving cytokine expression

Next, we sought to investigate how much of the variance in cytokine expression could be explained by biological and AD-related traits (**Figure 3**; **Supplementary Figure 3**). For this analysis, we used a linear mixed model to partition the variance attributable to multiple variables in the data (see Methods). The analysis included the same AD traits previously described, age at death, sex, a binary variable indicating whether the participant had at least one allele 4 harboring the *APOE* gene, and a measure of cerebral amyloid angiopathy (**Figure 3**; **Supplementary Figure 3**). With the bulk RNA-Seq data, we found that age at death was the strongest driver of variance in *C3* expression levels, accounting for 3.29% of the total variance. Cognitive decline was the next strongest factor, explaining 2.87% of the variance in *IL15* levels in bulk data. The global deposition of neurotoxic proteins had a 0.1% influence, while the *APOE* ε4 allele did not have a quantified influence on *IL15* expression (**Figure 3a** and **Supplementary Table 8**).

**Fig. 3:**
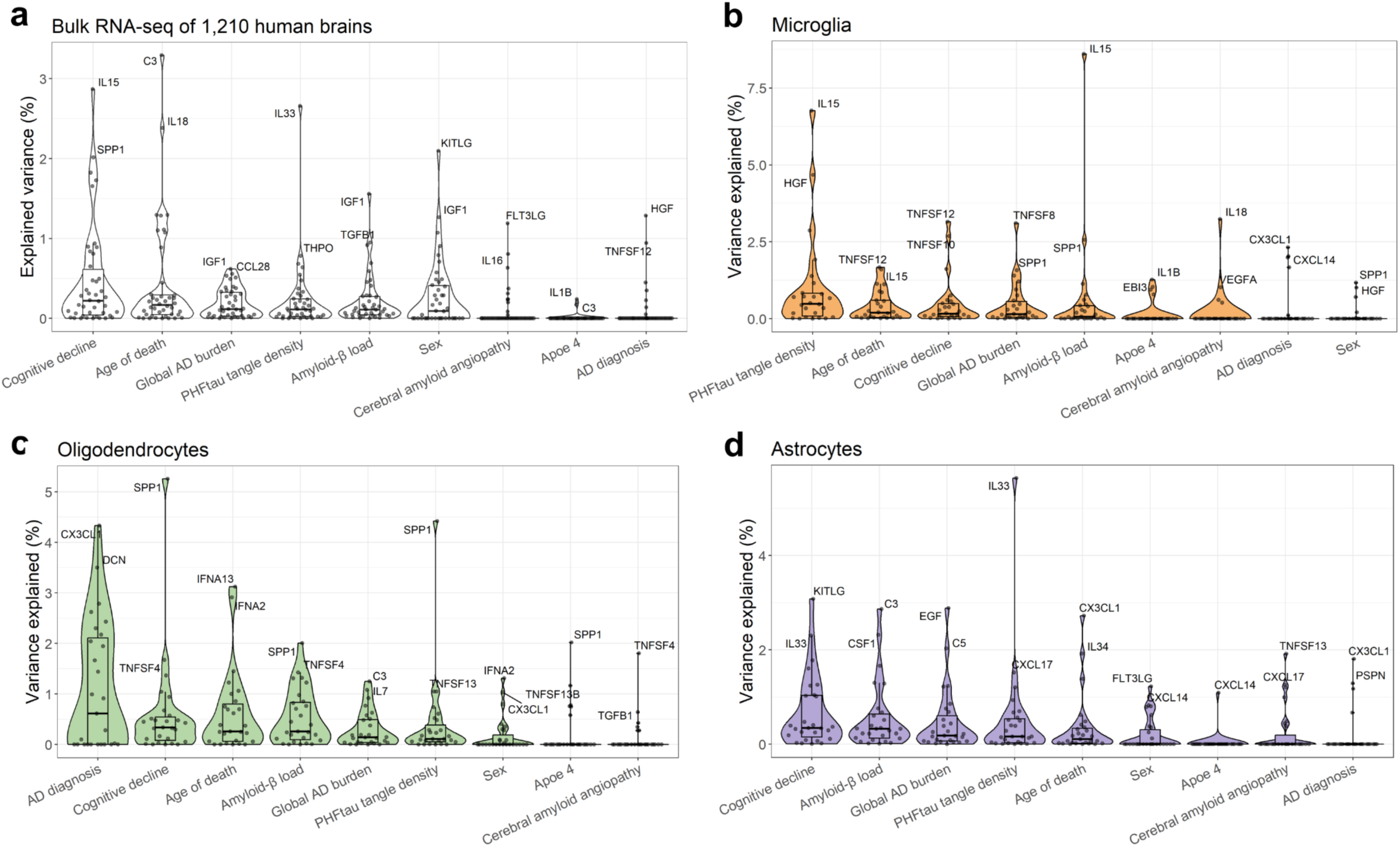
Contribution of biological drivers to cytokine expression variation. Distribution of variance explained per cytokine expression for **(a)** bulk RNA-Seq data from 1,210 participants and **(b, c, d)** from microglia, oligodendrocytes, and astrocytes, respectively. The y-axis represents the percentage of variance explained, while the x-axis shows the variance factors, with medians indicated by horizontal lines. The box spans the first to third quartiles, and the whiskers extend 1.5 times the interquartile range from the box. Each dot represents one gene, and the top two genes were labeled for each variable tested.

Overall, the max variance explained was between 3% in endothelial cells, 4% and 6% in neurons, and over 8% for the cytokines expressed in microglial cells (**Supplementary Figure 3**). For example, in microglia, amyloid-β load (8.6%) and PHFtau tangle density (6.76%) contributed most to the variance of *IL15* expression levels (**Figure 3b** and **Supplementary Table 9**). In oligodendrocytes, cognitive decline (5.25%), PHFtau tangle density (4.41%), and *APOE* ε4 allele (2.02%) explained part of the total variance in *SPP1* levels (**Figure 3c**). Amyloid-β load was the strongest driver of *IL7* variance in excitatory neurons (4.96%), inhibitory neurons (4.07%), and OPCs (5.06%) (**Supplementary Figures 3a–c**). PHFtau tangle density contributed to *IL33* variance in both bulk (2.65%) and astrocytes (5.63%), as shown in **Figures 3a** and **3d**, respectively. These results show that the percentage of variance explained for cytokines is cell-type dependent. Furthermore, AD-related traits explained a small fraction of the variance in cytokine expression, with cognitive decline, amyloid-β load, and PHFtau tangle density being the most significant contributors.

### AD genetic risk is associated with cytokine brain expression in specific cell types

To investigate the influence of genetic variants on cytokine expression in the cortex, we tested the association between cytokine expression and previously calculated polygenic risk scores for AD (AD-PRS) (Tasaki et al. 2019). First, we tested the association between cytokine expression and AD-PRS using linear regression (**Supplementary Table 10**). The top five significant findings from the association analysis are shown in **Figure 4a**. We observed that *C5* (*p*-value = 0.01538; t-value = −2.433369) and *IL33* (*p*-value = 0.01689; t-value = −2.398712) levels in astrocytes and *CCL28* (*p*-value = 0.01089; t-value = −2.557579) levels in excitatory neurons were inversely associated with higher AD risk (**Figure 4a**). Conversely, *HGF* levels in excitatory neurons (*p*-value = 0.0212; t-value = 3.091827) and *SPP1* expression in microglia (*p*-value = 0.00992; t-value = 2.590563) were positively associated with higher risk scores (**Figure 4a**).

**Fig. 4:**
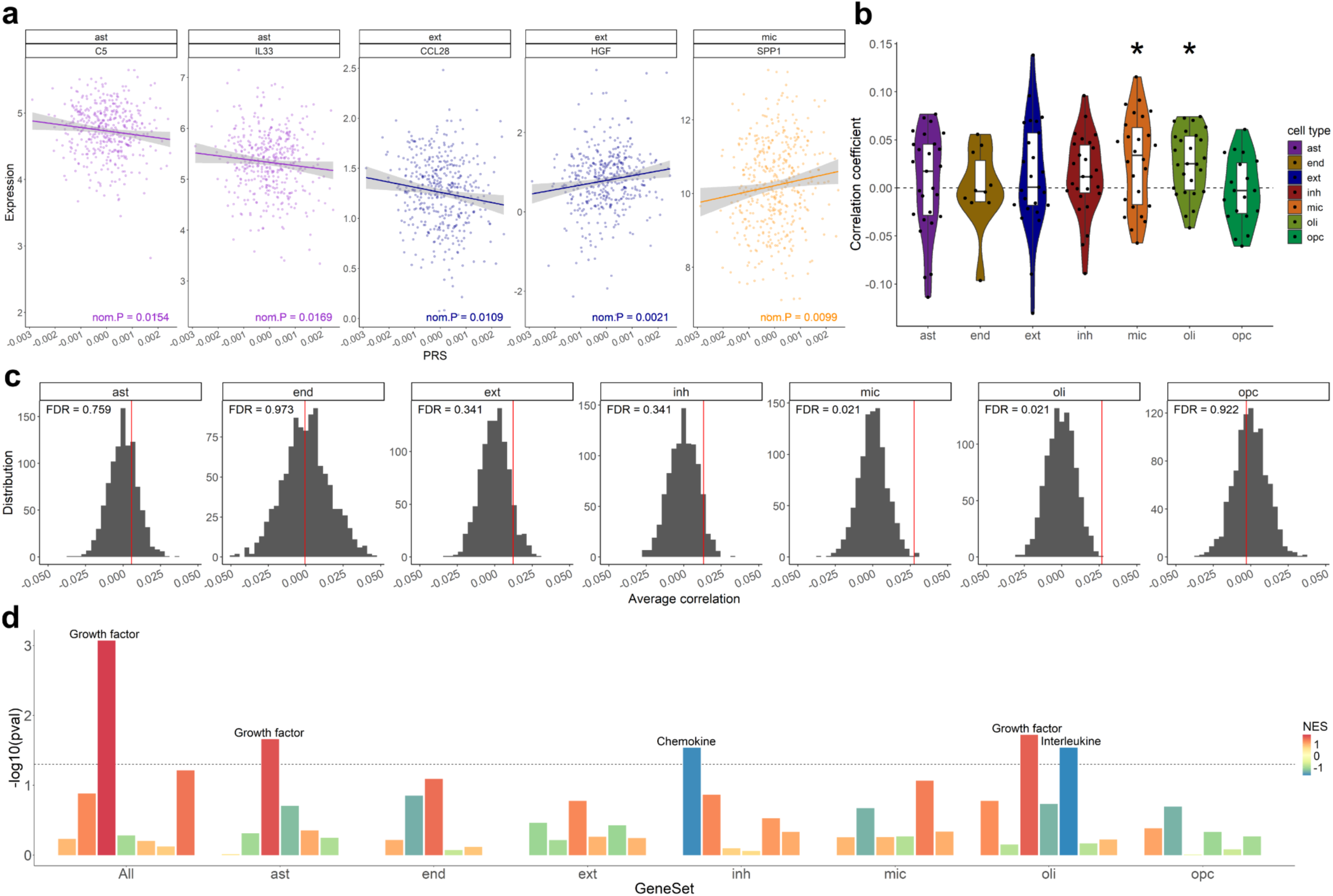
Cytokine production in distinct cell types was associated with the genetic risk score for Alzheimer’s disease. **a)** Scatter plot showing the correlation of the five cytokines with the most significant nominal *p*-values and the AD genetic risk score. **b)** Distribution of Spearman correlation coefficients between cell-type-specific cytokine expression and genetic risk score for Alzheimer’s disease in 424 individuals. Significant differences in mean correlation between cell-type-specific cytokines are indicated (\**p*-value < 0.05). Center lines represent medians, box limits show the 25th and 75th percentiles, whiskers extend 1.5 times the interquartile range, and outliers are shown as dots beyond the whiskers. The violin plots are color-coded by cell type. **c)** Distribution of mean correlation between AD-PRS and cytokine expression across cell types for 1,000 permutations, with the measured estimate indicated by a red line. Cytokine production in microglia and oligodendrocytes showed a significant association with AD-PRS (FDR *p*-value < 0.05). **d)** Bar plot of significantly enriched cytokine families associated with AD-genetic risk in each cell cluster from snRNA-Seq data. The y-axis shows the -log_10_ adjusted *p*-value, and the x-axis represents cell types. Colors represent the normalized enrichment score values from the gene set enrichment analysis, with red indicating positive values. PRS = polygenic risk score, ext = excitatory neurons, inh = inhibitory neurons, mic = microglia, ast = astrocytes, oli = oligodendrocytes, opc = oligodendrocyte progenitor cells, end = endothelial cells.

Participants with elevated AD risk scores were significantly associated with higher levels of microglia- and oligodendrocyte-derived cytokines (**Figure 4b**). A permutation test confirmed this association, which showed that higher AD risk was related to elevated cytokine expression in both cell types (**Figure 4c**). Even though the mean correlation between AD risk and inflammatory gene expression was higher in astrocytes and inhibitory neurons, it was not statistically significant (**Figure 4b**). The distribution of permuted mean correlations supported this pattern, showing no significant differences from the measured mean estimates in other cell types (**Figure 4c**).

Because cytokines can act together and be classified into families, we aimed to identify which families associated with AD-genetic risk are enriched in specific brain cell types. We conducted gene set enrichment analysis where the cytokine genes were ranked by family, using the t-statistics from the regression analysis of cytokine expression and AD-PRS (**Supplementary Table 11**). Among the genes expressed by all cells, the growth factor pathway was the most significantly enriched (*p*-value = 8.46 × 10^−4^; **Figure 4d**). This pathway was also significantly enriched in astrocytes (*p*-value = 0.022) and oligodendrocytes (*p*-value = 0.019). Using linear or logistic regression models, we found that the average expression of the growth factor family was associated with cognitive decline in bulk RNA-Seq data (adj *p*-value = 0.003; **Supplementary Figure 4a**), strongly associated with cognitive decline, amyloid-β, global AD burden, and AD diagnosis in astrocytes (adj *p*-value ≤ 0.001; **Supplementary Figure 4e**) and PHFtau tangle in endothelial cells (adj *p*-value = 0.005; **Supplementary Figure 4h**). The gene set comprising the interleukine family exhibited negative enrichment in oligodendrocytes (*p*-value = 0.029; **Figure 4d**), while the chemokine family demonstrated negative enrichment in inhibitory neurons (*p*-value = 0.029; **Figure 4d**). This indicates that their expression levels are reduced in association with AD-PRS within these specific cell types. No statistically significant association was observed between the interleukin family and AD traits in oligodendrocytes (**Supplementary Figure 4f)**; however, strong associations were found with amyloid-β load in neurons (adj *p*-value ≤ 0.001; **Supplementary Figures 4b–c**), and with cognitive decline, PHFtau tangles, and global AD burden in astrocytes (adj *p*-value ≤ 0.01; **Supplementary Figure 4e**). Additionally, the chemokine family was significantly associated with AD phenotype only in bulk RNA-Seq data, specifically with cognitive decline (adj *p*-value = 0.003; **Supplementary Figure 4a**). Together, these results offer valuable insights into the genetic influence of AD risk on brain inflammatory gene expression, deepening our understanding of neuroinflammatory mechanisms involved with Alzheimer’s dementia.

### Microglial cell subpopulation-specific cytokine signaling

Following up on our previous results, we investigated the myeloid cells of the brain and microglia. These cells had been previously characterized and annotated into 16 microglia subtypes (Green et al. 2024; Fujita et al. 2024), making them suitable for a systems-level analysis of cell communication. We analyzed intercellular communication networks within these subpopulations by predicting major signaling inputs and outputs using the CellChat pipeline (Jin et al. 2021). Full summary statistics are available in **Supplementary Tables 12-13**.

Focusing on cytokines, we identified 10 significant ligand-receptor pairs categorized into three key signaling pathway networks: SPP1, TGFβ, and Complement. In the inferred cell-cell SPP1 signaling network, we observed that lipid-associated Mic.13 cells were the dominant source of SPP1 ligands, followed by Mic.15, which is enriched with inflammation, stress signature genes, and disease-associated microglia TREM2-dependent markers (**Figure 5a-b**). The SPP1 signaling network in microglia was primarily mediated by paracrine signaling; however, Mic.13 (enriched in lipid signatures) and Mic.15 exhibited significant autocrine signaling (**Figure 5a**). Additionally, in this pathway, Mic. 15 acts as an important receiver and mediator of SPP1 signaling (**Figure 5b**). The SPP1-(*ITGAV*+*ITGB1*) ligand-receptor three-way pair was also identified as a potentially key mediator of the microglial communication network, with SPP1-(*ITGAV*+*ITGB5*) and SPP1-(*ITGA9*+*ITGB1*) contributing as well (**Figure 5c-d**). These results indicate that Mic.15 (enriched in inflammatory signatures) may play a pivotal role in the SPP1 signaling pathway, orchestrating and influencing communication among microglial subpopulations in the aged human brain (**Figure 5b**).

**Fig. 5:**
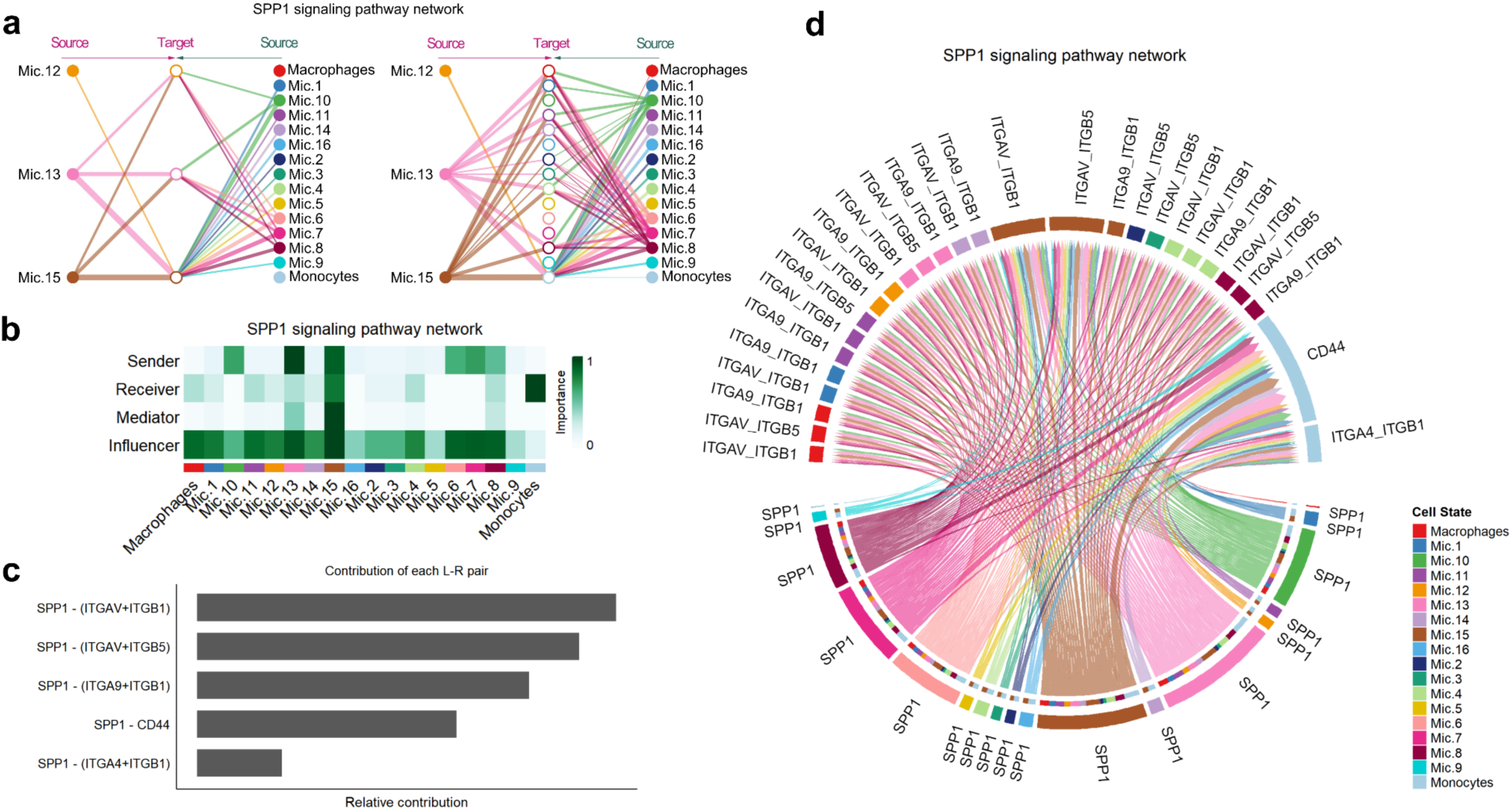
The cell-cell interactions among microglial subpopulations in the SPP1 signaling network. **(a)** Hierarchical plot shows interactions between microglial subpopulations in the inferred SPP1 signaling network. Solid circles indicate sources, while open circles indicate targets within the network. Intercellular interactions are represented by lines, with colors corresponding to the subtype source. The width of the edges reflects the communication probability. **b)** Heatmap shows the relative importance of each microglial cluster in the SPP1 signaling network, based on the computed four network centrality measures: sender, receiver, mediator, and influencer. **c)** Relative contribution of each ligand-receptor pair to the inferred signaling network. **d)** Chord diagram showing significant interactions among microglial subpopulations within the SPP1 signaling network. Inner bar colors represent the target cells, while outer bars correspond to the color-coded microglial sources.

In the inferred TGFβ signaling network, Mic.13 emerged as a major contributor in cell-cell communication. We noted that Mic.13 exhibits high target promiscuity, making it one of the primary targets within this network (**Supplementary Figures 5a-d; Supplementary Table 12**). By contrast, network centrality analysis showed that Mic.13 primarily acted as a stronger influencer in the inferred Complement signaling network, which is mediated by C3 levels and *ITGAX*-*ITGB2* receptors (**Supplementary Figures 5e-h; Supplementary Table 12**). The Mic.12, associated with human amyloid-β deposition (Gerrits et al. 2021), had moderate involvement in cell-cell communication, acting mostly as an influencer in the three signaling pathway networks (**Figure 5b; Supplementary Figures 5b-f**). These findings highlight the distinct roles of cytokines in microglial subpopulations in shaping or participating in intercellular communication networks in the aged human brain, with Mic.13 and Mic.15 emerging as key regulators within the SPP1, TGFβ, and Complement signaling pathways.

## DISCUSSION

In this study, we examined cytokine gene expression in aged human brains using bulk and single-nucleus RNA sequencing (snRNA-Seq) from the ROSMAP cohorts. By applying linear and/or logistic regressions, we identify specific associations between cytokine expression levels and AD-related traits. Next, we noted that these AD traits contribute differently to expression variation, with amyloid accumulation driving the highest variance in microglial cells when focusing on cytokines. Our analysis also showed that AD genetic risk was associated with cytokine production, as high-risk individuals exhibited increased cytokine levels from microglia and oligodendrocytes. Finally, cell communication analysis revealed that Aβ-associated (Mic.12/13) and inflammation-related (Mic.15) microglia may regulate SPP1, TGFβ, and Complement signaling pathways among microglial cells in the aged human brain.

Neuroinflammation has been suggested to play an independent role from the early stages of AD pathology, with pro- and anti-inflammatory cytokines acting together. However, their chronic pathological expression may disrupt homeostasis, leading to neuronal damage and cognitive deficits (Chen et al. 2024; Bourgognon and Cavanagh 2020). Studies suggest that elevated pro-inflammatory *IL-15* in serum and cerebrospinal fluid is associated with the severity of cognitive decline and increased levels of total and phosphorylated tau (Bishnoi, Palmer, and Royall 2015; Rentzos et al. 2006; Taipa et al. 2019; Janelidze et al. 2018; Popp et al. 2017; Daniilidou et al. 2023; Rentzos and Rombos 2012). This evidence indicates a significant role of *IL-15* in the mechanisms underlying AD, highlighting its potential as a biomarker. Although cytokines play a vital role in brain homeostasis, their expression in cortical tissue remains largely unexplored in human-aged brains. Also, the neuroinflammatory complexity and limited temporal-spatial resolution make it difficult to define glial roles in AD progression and their involvement in histopathological changes like protein deposition and neuronal loss (Vidovic and Spittau 2024). A study of 81 brain tissue samples from ROSMAP volunteers found that individuals with Aβ deposition had significantly higher concentrations of specific inflammatory cytokines in cortical tissue (Flores-Aguilar et al. 2021). More recently, *SPP1* overexpression within the brain parenchyma has been linked with a higher risk of severe neurodegenerative diseases, including AD. Furthermore, its expression in specific microglial subgroups (homeostatic/tau-associated Mic.3 and lipid-associated Mic.12 and 13) correlates with increased cognitive decline (Lopes et al. 2024). While these findings provide valuable insights into cytokine activity in Alzheimer’s dementia, our study builds on this knowledge by utilizing a larger sample size and investigating a broader array of cytokine genes expressed by brain cell types.

Here, we evaluated cytokine gene expression profiles across all major cortical brain cell types and compared the differential expression patterns among individuals based on their clinical diagnoses and protein deposition. As expected, the associations between cytokines and AD traits differed across cell types, with some significant signals being captured only in the snRNA-Seq. These results suggest that, at the cellular level, cytokine expression patterns may vary among cells in the brains of older adults. We also found that age at death was the major contributor to the variance in cytokine expression in the bulk data. When examining specific cell groups, protein deposition and cognitive decline in cortical tissue emerged as important drivers of variance in cytokine expression. Another source of variation in cytokine gene expression was the genetic risk for AD. Further supporting the link between AD genetic risk and cytokine expression, we observed that a higher AD-PRS score correlated with increased *SPP1* expression in microglia and decreased *IL33* expression in astrocytes. Microglial subpopulations Mic.12, Mic.13, and Mic.15 (enriched with inflammatory expressed genes), identified as being linked to pathological AD and neuroinflammation, have a significant influence on *SPP1* signaling in aged brain samples, suggesting its involvement in the molecular mechanisms underlying cognitive decline and protein deposition.

This study has strengths and limitations. A key strength is the integration of bulk and single-nuclei analyses, which provides a comprehensive view of cytokine expression by distinct brain cell types in the human brain. The large sample size and standardized annual clinical assessments, aligned with high follow-up and brain autopsy rates, ensure a well-characterized and deeply phenotyped cohort, enhancing the association between cytokine expression and AD traits. ROSMAP consists of community-based studies where participants were free of dementia at enrollment, minimizing referral bias. However, these cohorts are characterized by highly educated individuals with European ancestry. Therefore, these findings cannot be extrapolated to individuals with different social and genetic backgrounds. Further research is needed to better understand cytokine expression patterns across brain regions beyond the DLPFC. Under homeostatic conditions, cytokines play a key role in cognitive function, particularly in hippocampal-dependent learning and memory (Bourgognon and Cavanagh 2020). Given that hippocampal neurodegeneration is a key hallmark of AD (Rao et al. 2022; Pini et al. 2016), exploring this region may reveal novel or stronger associations between cytokines and AD traits. Additionally, the high mean age of volunteers may have influenced these results. In future investigations, it might be possible to use a younger cohort and incorporate a broader range of inflammatory molecules, including non-coding RNAs such as miRNAs, piRNAs, and circRNAs.

## METHODS

### Study cohorts: participants and clinical-neuropathological evaluations

Data were obtained from two longitudinal clinical-pathologic cohort studies focused on risk factors for cognitive decline and Alzheimer’s disease (AD). The Religious Order Study (ROS) and the Rush Memory and Aging Project (MAP), collectively called ROSMAP, were launched in 1994 and 1997 (David A. Bennett et al. 2018). At enrollment, all participants were without known dementia and consented to detailed annual cognitive and clinical assessments. They signed the Anatomical Gift Act for brain donation and provided informed and repository consents. An Institutional Review Board at Rush University Medical Center approved both studies. The phenotypic variables included in this manuscript were defined as follows:

#### Global AD pathology burden, amyloid-β load, and PHFtau tangle density

To quantify amyloid-β deposition and PHFtau tangle density, brain autopsies were conducted by trained staff blinded to age and clinical data. The brains were removed, weighed, and fixed in 4% paraformaldehyde for neuropathological evaluation. Tissue samples were collected from eight distinct brain regions: the angular gyrus, anterior cingulate, calcarine cortex, entorhinal cortex, hippocampus, inferior temporal cortex, midfrontal gyrus, and superior frontal cortex. Immunohistochemical staining with Aβ and tau antibodies on 20-micron sections, combined with image analysis and stereology, was employed to assess the neuropathological burden (Kapasi et al. 2023). A square root transformation was applied to the data for analysis. For unbiased assessments, examiners were blinded to clinical data during all evaluations. Global AD pathology burden quantifies Alzheimer’s pathology by aggregating counts of neuritic plaques, diffuse plaques, and neurofibrillary tangles through microscopic examination of silver-stained slides from five regions: midfrontal cortex, mid-temporal cortex, inferior parietal cortex, entorhinal cortex, and hippocampus (D. A. Bennett et al. 2003).

#### Cognitive decline and resilience

ROSMAP participants completed a series of 21 cognitive performance tests each year, administered by blinded examiners unaware of their earlier performance. The rate of global cognitive decline reflects individual changes in cognition over time. The cognitive examination included 21 tests, 17 of which were used to create a global composite measure of cognitive function. Raw scores for individual tests were standardized using the baseline means and standard deviations of the entire cohort and then averaged across tests to obtain the composite score. This rate was estimated using a linear mixed-effects model, with global cognition as the longitudinal outcome and adjustments made for age at baseline, sex, and years of education (Wilson et al. 2015). To calculate the cognitive resilience variable, linear mixed-effects models measured the slope of global cognition adjusted for the same demographic variables plus all measured neuropathology indices (Boyle et al. 2021; Lei Yu et al. 2020).

#### Alzheimer’s dementia and MCI diagnoses

A clinical diagnosis of cognitive status is made by a neurologist with expertise in dementia and a clinician who reviews all clinical data available, both remaining blinded to patient demographic information. A detailed assessment of available clinical data was performed to determine the most likely clinical diagnosis at the time of death, ensuring that the reviewers could not access post-mortem data (Schneider et al. 2007; David A. Bennett et al. 2006). Diagnosis of MCI and clinical Alzheimer’s dementia follows the criteria set by the National Institute of Neurological and Communicative Disorders and Stroke and the Alzheimer’s Disease and Related Disorders Association (NINCDS/ADRDA). Mild cognitive impairment (MCI) is diagnosed when cognitive impairment is present but does not meet the criteria for dementia. Individuals without dementia or MCI are classified as having no cognitive impairment (NCI) (David A. Bennett et al. 2006; D. A. Bennett et al. 2002).

### Cytokine gene list

To investigate whether abnormal deposition of pathological proteins and cognitive decline were linked with neuroinflammation, brain cytokine production was assessed using a panel of 147 genes (**Supplementary Table 14**). A curated list of 147 cytokine genes was generated based on supplementary data from two key studies. These studies provided a comprehensive view of cytokine regulation and immune responses, mapping transcriptional control networks and detailing cell-type-specific cytokine activity at single-cell resolution (Cui et al. 2023; Santoso et al. 2020). This list was used as the basis for the subsequent analysis of cytokine gene expression in this manuscript. After filtering bulk and single-nuclei data to include only cytokine genes, we evaluate their association with AD phenotype through linear or logistic regressions. Since age, sex, and years of education are important sources of confounders, we adjusted our statistical model to account for them, as in previous ROSMAP publications (Lei Yu et al. 2018; L. Yu et al. 2023).

### Bulk RNA-seq data: collection and analysis

We utilized previously generated and processed RNA-Seq data from DLPFC tissue, as described in Mostafavi et al. and Tasaki et al. (Mostafavi et al. 2018; Tasaki et al. 2022). Briefly, the data were obtained using Illumina platforms, aligned to the human genome (GENCODE Release 27 GRCh38) using STAR v2.6, and transcript-level quantification was performed with Kallisto v0.46. The dataset included 17,294 genes detected in over 50% of the samples. Prior normalization procedures included Conditional Quantile Normalization, log₂-transformation of counts per million (CPM), and adjustment for technical confounders. Technical confounders were accounted for using linear regression models that included post-mortem interval, sequencing batch, RNA quality number (RQN), total spliced reads, and metrics derived from Picard and Kallisto.

### Single-nuclei RNA-seq data: collection and analysis

Nuclei isolation was originally conducted in 479 brain specimens using available frozen pathological material from the DLPFC, processed via dounce homogenization and centrifugation (Green et al. 2024; Fujita et al. 2024). Single-nuclei profile was performed on pooled sample batches to boost library throughput and reduce batch effects and costs. Each library construction batch included eight participants, resulting in 60 batches. Even though samples were randomly assigned, they were balanced by clinical, pathological diagnoses, and sex. In each batch, the nuclei suspension from eight donors was combined. Single-nucleus RNA-Seq libraries were prepared using the 10x Genomics 3 Gene Expression kit (v3 chemistry), following the manufacturer’s protocol. After sequencing, 127 pooled libraries were created. Alignment, noise removal, demultiplexing, normalization, and classification were performed. Low-quality nuclei and doublets were removed using cell-type-specific thresholds. The preprocessing and quality control (QC) steps were extensively described previously (Green et al. 2024). In brief, approximately 1.64 million high-quality nuclei profiles were retained from 465 individuals. ElasticNet-regularized logistic regression was performed to annotate cells, resulting in seven major cell types. Our analysis was conducted for all the major cell types: astrocytes (ast), excitatory neurons (ext), inhibitory neurons (inh), microglia (mic), oligodendrocytes (oli), oligodendrocyte precursor cells (OPCs), and endothelial cells (end). In each cell type, the expression profile of marker genes was identified and categorized using the Seurat *FindAllMarkers* function. The sub-clustering analysis resulted in 96 subpopulations. After preprocessing and QC analyses, 424 participants were retained for the expression analysis (Lopes et al. 2024; Fujita et al. 2024).

Pseudo-bulk matrices were created by summing counts per cell for each participant. We retained genes with at least one count per million (CPM) in 80% of the samples, and TMM was applied with voom for normalization (Law et al. 2014).

### Generalization cohorts

For generalization analyses, we selected two external bulk RNA-seq datasets: the Mayo Clinic and the Mount Sinai Brain Bank (MSBB). The Mayo Clinic dataset includes 597 brain tissue samples isolated from the temporal cortex and cerebellum. Transcriptomic data correspond to 355 unique individuals from the Mayo Brain Bank and the Banner Sun Health Institute (Allen et al. 2016). The MSBB dataset contains RNA-seq data from 1,282 samples from multiple brain regions, covering 316 unique individuals (Wang et al. 2018). Both datasets were uniformly processed through the AMP-AD RNA-Seq Harmonization Study (syn21241740). Matrices with conditional quantile normalized (CQN) expression values were obtained for the subsequent analysis (Mayo: syn27024965, and MSBB: syn27068756). We filtered the datasets to include only samples functionally connected to the DLPFC (i.e., 258 temporal cortex samples from Mayo and 304 frontal cortex samples from MSBB). Both datasets were further filtered to include only targeted cytokine genes, and association analyses were performed with the available AD traits. We tested for associations with diagnosis, Braak stage, and Thal score in the Mayo Clinic dataset. In the MSBB dataset, the available variables were CERAD score, Braak stage, Clinical Dementia Rating (CDR), and neuritic plaque density.

### Statistical methods

To evaluate the association between cytokine expression and AD traits, we used a linear regression model for continuous outcomes and logistic regression for categorical ones. Both models were adjusted for age, sex, and years of education to control for potential confounders. We conducted Pearson’s and Spearman’s correlation analyses to examine variable relationships. The differential expression of inflammatory mediators in individuals diagnosed with AD, MCI, and controls was assessed using two-tailed Student’s t-tests for two-group comparisons. Data were presented in boxplots, with the median indicated by a horizontal line and whiskers representing the 25th and 75th percentiles.

#### VariancePartition

To identify and quantify sources of biological variation in cytokine expression, we applied the variancePartition function in R. This package applies linear mixed models to quantify the influence of biological and technical factors on gene expression, revealing patterns of variation across datasets (Hoffman and Schadt 2016).

#### Genetics

Next, we examined the influence of AD genetic risk on cytokine expression using the polygenic risk scores (AD-PRS) previously calculated by (Tasaki et al. 2019) for all 424 individuals in the single-nuclei RNA-seq data. Using Spearman correlation, we assessed the relationship between AD-PRS and the average cytokine expression in each cell cluster. Permutation tests were applied to assess the statistical significance of the associations in each cell type. For this purpose, we followed the phenotype permutation shuffling strategy as implemented in GSEA (Subramanian et al. 2005), which preserves the correlations between genes in the dataset across permutations. Null distributions were obtained by taking 1,000 average correlations between AD-PRS and randomly shuffled cell-specific cytokine expressions. *P*-values were calculated by comparing the observed correlation with the null distribution for each cell type. Benjamini–Hochberg was used to adjust *p*-values to test multiple cell types.

Finally, we performed the PRS analysis by cytokine families. First, we calculated the mean expression for each annotated family. Then, we performed gene set enrichment analysis (GSEA) using the cytokine families list as the gene set. The cytokine genes were ranked based on t-values derived from the correlation analysis.

### Cell communication

Cell–cell communication analysis was conducted using CellChat v2, an R package designed for inferring, analyzing, and visualizing intercellular interactions from snRNA-seq data (Jin, Plikus, and Nie 2025). Only microglial subpopulations annotated from snRNA-seq were included in the analysis. By detecting overexpressed ligands and receptors, CellChat predicts significant communications and quantifies them with probability values. Ligand–receptor pairs with a *p*-value of less than 0.05, determined by CellChat, were considered significant interacting molecules between microglial subpopulations. To focus on cytokine signaling pathways, we further filtered the significant associations to include only those involving cytokine genes. Our analysis followed the official step-by-step protocol. All cell communication graphs were constructed utilizing the built-in functions of the CellChat package.

## DATA AVAILABILITY

Transcriptomic data used in this manuscript are available with the following accession numbers: ROSMAP bulk DLPFC (syn3388564) and snRNA-Seq (syn31512863). The phenotype data can be requested at the RADC Resource Sharing Hub at www.radc.rush.edu or the AD Knowledge Portal (https://adknowledgeportal.org).

## CODE AVAILABILITY

The relevant code generated by this study is available on the GitHub repository https://github.com/RushAlz/Cytokines_public_release.

## ACKNOWLEDGEMENTS

We thank all the ROSMAP participants and the investigators, and the staff at the Rush Alzheimer’s Disease Center. ROSMAP is supported by the following grants: P30AG10161, P30AG72975, R01AG15819, R01AG17917, U01AG46152, and U01AG61356. Student fellowship from the Brazilian agency CAPES (Coordenação de Aperfeiçoamento de Pessoal de Nível Superior).

## AUTHOR CONTRIBUTIONS

K.P.L supervised the data analysis; K.P.L and D.G conceived and supervised the study; J.B.G curated the cytokine list, ran the regression analysis for both bulk and single-nucleus data, performed exploratory analysis, and created the github pages; R.A.V developed regression functions for the analysis and conducted the genetic association study; L.S ran the cell-communication pipeline; R.T.R supervised the cell communication analysis; Y.W sequenced the bulk RNA-Seq samples; V.M and P.L.D.J provided the single-nucleus dataset; J.A.S supervised the brain pathology quantification; S.T ran the pipelines for bulk RNA-Seq quantification, including normalization and data adjustment; D.A.B built the ROS and MAP cohorts; J.B.G. and K.P.L. wrote the manuscript with feedback from the co-authors. All authors approved the manuscript.

## COMPETING INTERESTS

The authors declare no conflict of interest.

## Notes

https://github.com/RushAlz/Cytokines_public_release

### Competing Interest Statement

The authors have declared no competing interest.

