## Supplementary figures for "Cytokine expression profile in the human brain of older adults"

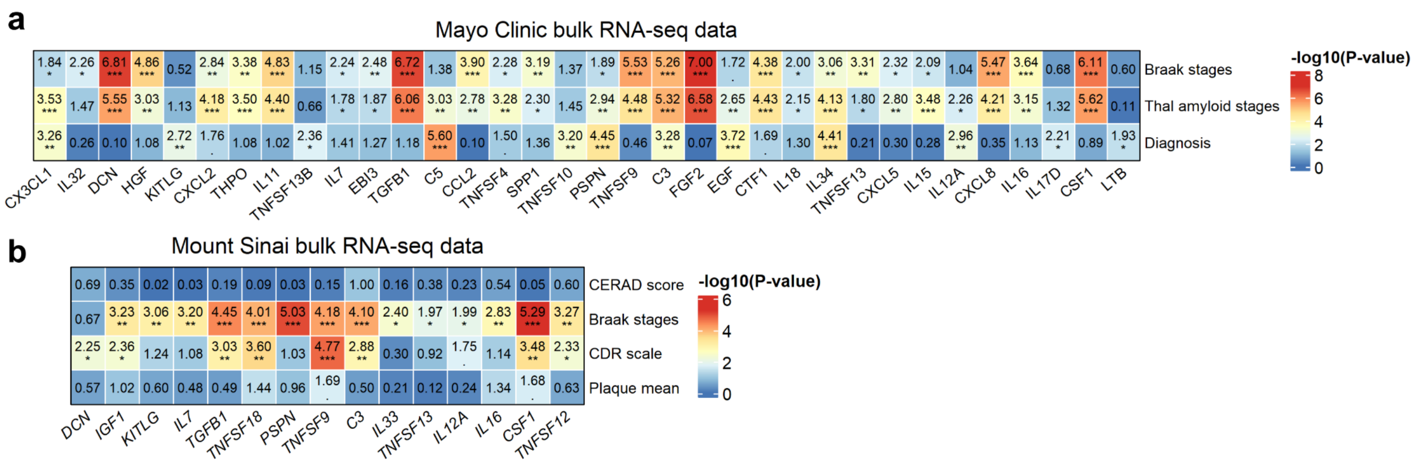


**Supplementary Fig. 1: Replication analysis using bulk RNASeq dataset from the Mayo Clinic and Mount Sinai datasets.** Heatmaps showing significant associations between cytokine expression levels and AD traits using **(a)** the Mayo Clinic dataset and **(b)** the Mount Sinai Brain Bank dataset. Linear and/or logistic regressions between cytokine expression (columns) and phenotypic data (rows). All models were adjusted for age, sex, and education, with the Mount Sinai data also adjusted for ethnicity. Numbers and colors represent the -$\log_{10}$ of the p-value (*** FDR < 0.001; ** FDR < 0.01; * FDR < 0.05; . FDR = 0.1), ranging from highest (red) to lowest (blue).

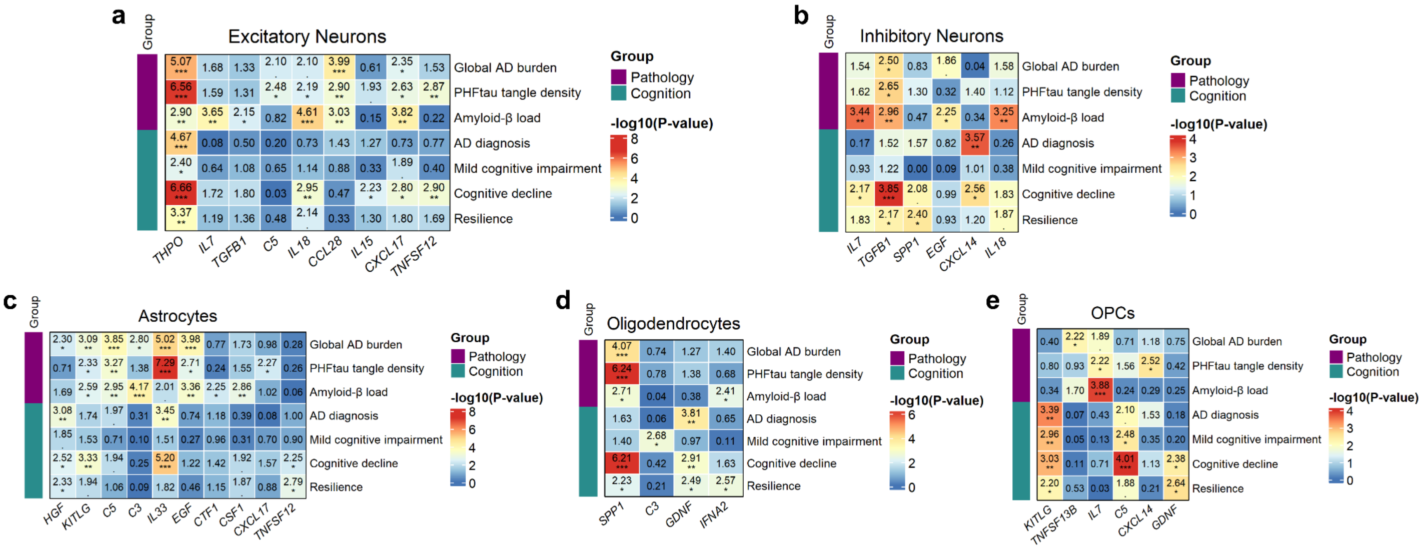


**Supplementary Fig. 2: Association analysis between cytokine expression and AD traits by cell type.** Heatmaps showing significant associations between cytokine expression levels and AD phenotypes across five distinct cell types: **(a)** excitatory neurons, **(b)** inhibitory neurons, **(c)** astrocytes, **(d)** oligodendrocytes, and **(e)** oligodendrocyte progenitor cells (OPC cells). Linear and/or logistic regressions were conducted to examine relationships between cytokine expression in the DLPFC region (columns) and phenotypic data (rows). All models were adjusted for age, sex, and education to account for confounding factors. The AD-traits are labelled according to their category: Pathology (global AD pathology burden, PHFtau tangle density, and amyloid-β load) and Cognition (AD clinical diagnosis, MCI diagnosis, cognitive decline, and resilience). Numbers and colors represent the -$\log_{10}$ of the p-value (*** $P < 0.001$, ** $P < 0.01$, * $P < 0.05$, . $P = 0.1$), ranging from highest (red) to lowest (blue).

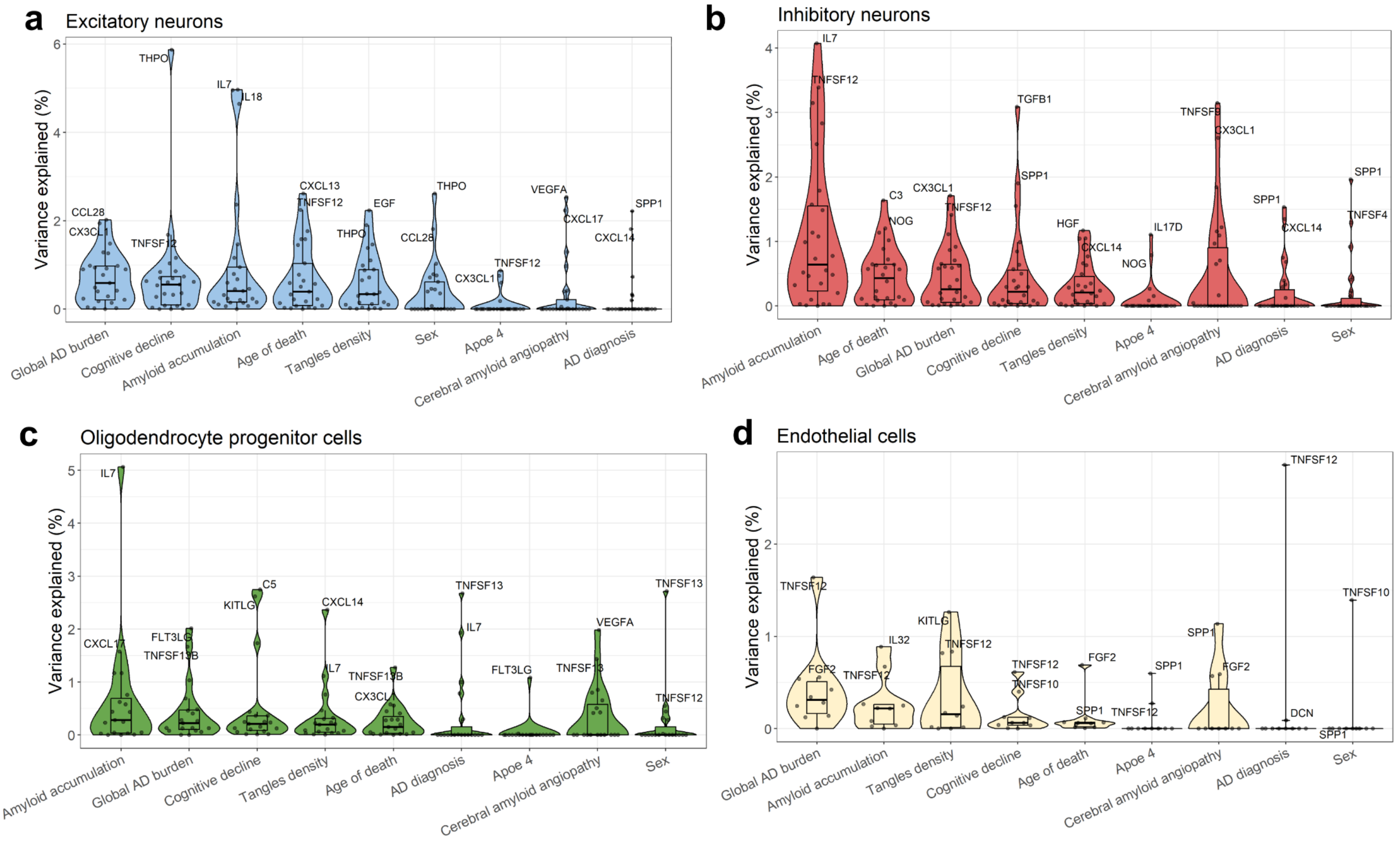


**Supplementary Fig. 3: Contribution of biological drivers to cytokine expression variation by cell type.** Variance partition analysis using AD phenotypic data to assess the percentage of explained variance in cytokine expression for **(a)** excitatory neurons, **(b)** inhibitory neurons, **(c)** oligodendrocyte progenitor cells (OPC cells), and **(d)** endothelial cells. The y-axis represents the percentage of variance explained, while the x-axis shows the variance factors, with medians indicated by horizontal lines in the box plots.

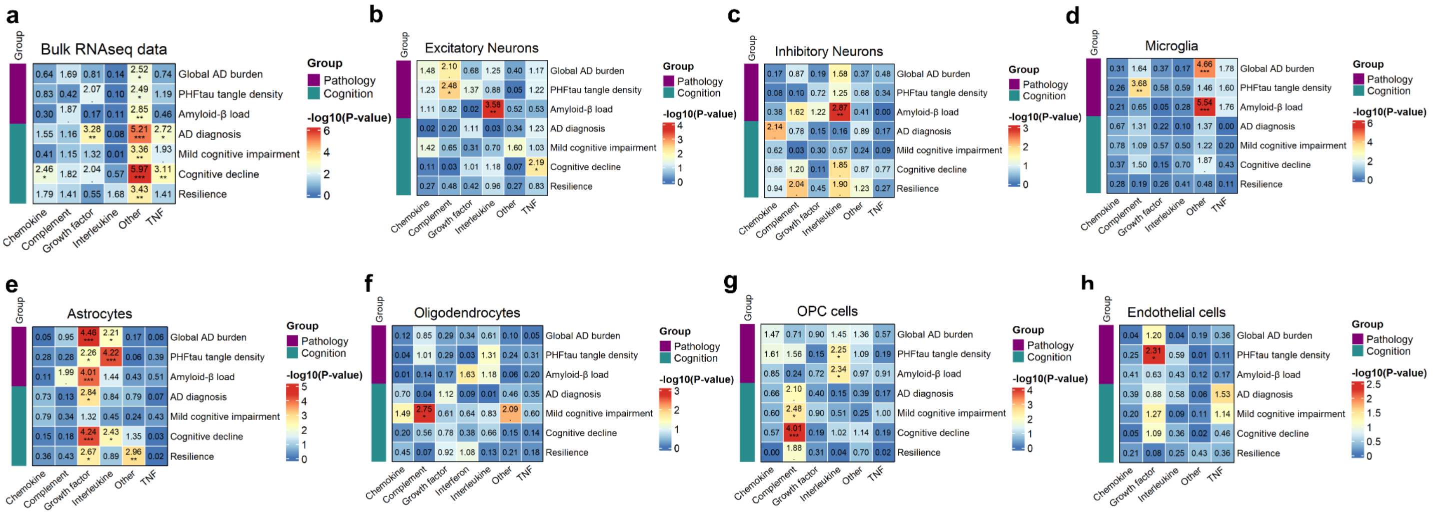


**Supplementary Fig. 4: Association analysis between average cytokine family expression and AD phenotype variables in cell clusters.** Heatmaps show significant associations between the average expression of cytokine families and AD phenotypic data in bulk data and across seven cell types in the single-nuclei dataset. Linear and/or logistic regressions were used to analyze the relationships between average cytokine family expression (columns) and phenotypic data (rows). The AD-traits are labelled according to their category: Pathology (global AD pathology burden, PHFtau tangle density, and amyloid-β load) and Cognition (AD clinical diagnosis, MCI diagnosis, cognitive decline, and resilience). Colors represent the -log10 of the p-value (*** P < 0.001, ** P < 0.01, * P < 0.05, . P = 0.1), ranging from highest (red) to lowest (blue).

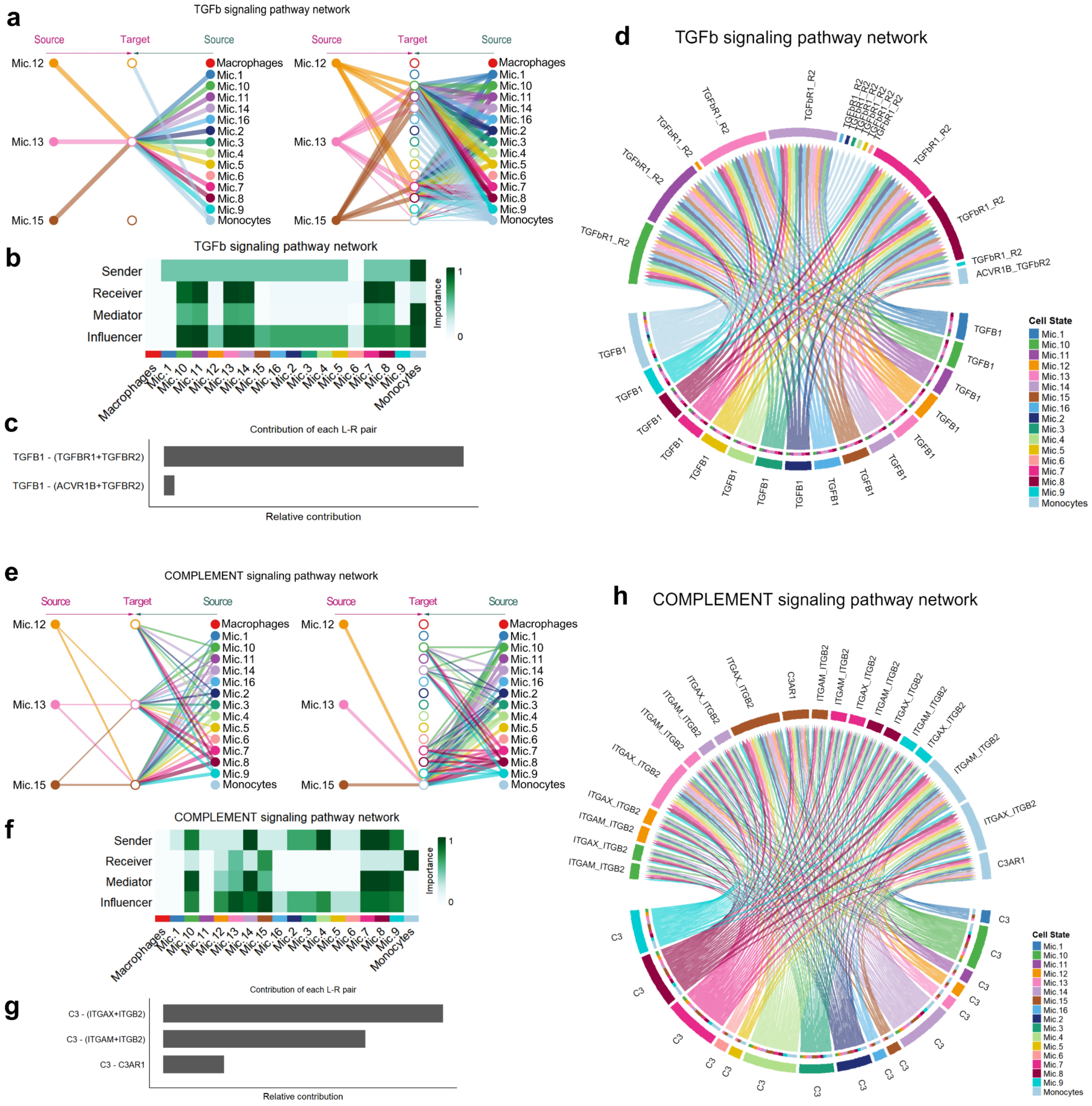


**Supplementary Fig. 5: The cell-cell interactions among microglial subpopulations in the TGFb and Complement signaling network. a)** Hierarchical plot showing interactions between microglial subpopulations in the inferred TGFb signaling network. Solid circles indicate sources, while open circles indicate targets within the network. Intercellular interactions are represented by lines and colors corresponding to the source. The width of the edges reflects the communication probability. **b)** The heatmap shows the relative importance of each microglial cluster in the TGFb signaling network based on the four computed network centrality measures: sender, receiver, mediator, and influencer. **c)** Relative contribution of each ligand-receptor pair to the inferred TGFb signaling network. **d)** Chord diagram showing significant interactions among microglial subpopulations within the TGFb signaling network. Inner bar colors represent the target cells, while outer bars correspond to the color-coded microglial sources. **e)** The CellChat-inferred Complement signaling network. **f)** Computed network centrality analysis showing the relative importance of microglial subpopulations in the complement signaling network. **g)** Relative contribution of each C3 ligand-receptor pair. **h)** Significant microglial interactions in the inferred complement signaling network.
